# Multi-specific nanobody-Fc fusions against TcdB protect *in vivo* from diverse *Clostridioides difficile* strains

**DOI:** 10.64898/2026.11.12.766007

**Authors:** Alyssa G. Ehni, Bruno B. C. Lança, Heather K. Kroh, Anna J. Smith, Audrey K. Thomas, M. Kay Washington, Benjamin W. Spiller, D. Borden Lacy

## Abstract

*Clostridioides difficile* infection (CDI) is an urgent antibiotic-associated threat and the leading cause of nosocomial diarrhea in the United States, with a high rate of recurrence. Antibiotics are the standard of care for treating CDI but can prolong microbial dysbiosis in the gut. The only FDA-approved monoclonal antibody (mAb) for the prevention of recurrent CDI, Zinplava™ (bezlotoxumab), was discontinued in January 2025. The target of bezlotoxumab is the dominant virulence factor of *C. difficile*, the exotoxin B (TcdB). However, TcdB sequences vary across strains and encompass twelve distinct subtypes. Consequently, sequence variation between subtypes presents a challenge for the identification of pan-neutralizing mAb therapeutics. Here, we describe the generation of three bi-specific and one tri-specific nanobody-Fc fusions targeting TcdB. Each fusion neutralizes TcdB *in vitro* across clinically relevant subtypes with greater potency than bezlotoxumab. In a mouse model of CDI, using three diverse *C. difficile* strains, several of the fusion proteins improve infection outcomes over bezlotoxumab when administered prophylactically. We also highlight one fusion that improves survival in a therapeutic mouse model of CDI. These data demonstrate multi-specific nanobody-Fc fusions as a strategy to improve potency and mitigate the impact of sequence variation within the toxin. Our fusions outperform the standards established by bezlotoxumab and represent strong candidates for translation.

**One sentence summary:** Bi- and tri-specific nanobody-Fc fusions confer broad TcdB neutralization and protect *in vivo* against three diverse strains of *C. difficile*.

## Introduction

*Clostridioides difficile* is a Gram-positive, spore-forming, anaerobic bacterium and is the causative agent of *C. difficile* infection (CDI) (*1*). CDI is an antibiotic-associated disease, as the depletion of gut microbial diversity provides *C. difficile* a colonization niche within the colon (*1*). The first line of defense against CDI is antibiotics, most commonly vancomycin or fidaxomicin (*2*). However, antibiotic treatments can prolong gut dysbiosis in patients and may contribute to the high incidence of recurrent CDI (*2*). Anywhere from 25-35% of patients experience recurrent CDI after the first infection, and the risk of developing further recurrent infections increases with each relapse (*2, 3*). The sole monoclonal antibody (mAb) approved for treatment of recurrent CDI, Zinplava™ (bezlotoxumab), was effective, but it was discontinued in January 2025 (*4, 5*). Alternative recurrent CDI treatments are limited to donor-derived fecal microbiota products (REBYOTA® and VOWST™) or traditional fecal microbiota transplantation (FMT) (*2, 6*), but these options have inherent cost, safety, and efficacy limitations.

While *C. difficile* strains can produce up to three different toxins, TcdA, TcdB, and the CDT binary toxin, the cellular damage of CDI is mainly mediated by toxin B (TcdB) (*1, 7*). TcdB is a large exotoxin (270 kDa) composed of four domains: the glucosyltransferase domain (GTD), autoprocessing domain (APD), delivery domain (DD), and the combined repetitive oligopeptides domain (CROPs) (*1*). Following receptor binding, TcdB enters host cells through receptor-mediated endocytosis. Subsequent endosome acidification triggers the DD to form a pore in the endosomal membrane, which translocates the APD and GTD into the cytoplasm. In the presence of inositol hexaphosphate, the APD releases the GTD, which glucosylates and inactivates Rho family GTPases (*1*). This disrupts the cytoskeleton, causes cytokine release, and eventually leads to cell death (*1*). During infection, the effects of TcdB manifest as a range of symptoms from diarrhea to pseudomembranous colitis and toxic megacolon (*1*). TcdB alone is sufficient to cause disease in animal models, and in Phase III clinical trials for bezlotoxumab, the addition of an anti-TcdA mAb did not improve efficacy over bezlotoxumab alone (*4, 7, 8*). Further, there are strains of *C. difficile* from the 017 ribotype that only produce TcdB and have been associated with outbreaks (*9*).

Since TcdB is the primary driver of cellular damage, it is a promising therapeutic target for CDI. However, TcdB sequences vary across strains, with at least twelve defined subtypes (*10*). TcdB1/2/3 are the most translationally relevant as they are found in the majority of clinical isolates (*10, 11*). Such sequence divergence complicates the development of a pan-neutralizing anti-TcdB therapeutic that targets a conserved site across subtypes. Bezlotoxumab is an example where it was developed from a mAb targeting the CROPs domain of TcdB1 (*12*). However, the CROPs domain is less conserved between TcdB1 and TcdB2, so consequently, bezlotoxumab lost some affinity for TcdB2 due to amino acid mutations in the bezlotoxumab epitope (*11, 13*).

Nanobodies (Nbs) are an emerging class of biologic drugs, with the first Nb-based drug approved in 2018 (*14*). Nbs are single variable heavy-chains derived from camelid or shark antibodies (*15*). Nbs offer great specificity and affinity and are manipulable due to their small, single-domain structure (15 kDa) (*15*). A limitation of using Nbs as therapeutics is their short half-lives, ranging from 1 – 2 hours owing to their molecular weight being below the threshold for glomerular filtration of approximately 50 kDa (*14*). The fusion of a Nb to the Fragment Crystallizable (Fc) portion of an antibody (Nb-Fc) is one method to increase the half-life by increasing the molecular weight above the glomerular filtration cutoff and by allowing recycling through the neonatal Fc receptor (FcRn) (*16*). Another advantage of a Nb-Fc fusion is an increase in valency (*17*). Nb-Fcs can be further engineered into bi-specific proteins, with a “knobs-into-holes” design as one approach that forces heterodimerization (*18–20*). Tri-specific fusions are also possible by fusing a Nb to a light-chain constant region and adding a CH1 domain to the Fc for pairing similar to that of a native antibody (*21*). We hypothesized that engineering anti-TcdB Nb-Fcs would generate a novel therapeutic capable of targeting multiple sites of TcdB to overcome subtype variations and improve neutralization potency. In this study, we demonstrate that engineered Nb human Fc fusions (Nb-hFc) targeting multiple TcdB epitopes located in the DD and GTD improve *in vitro* neutralization and protect *in vivo* against three diverse *C. difficile* strains with greater efficacy than the industry standard, bezlotoxumab.

## Results

### Multi-specific Nb-hFc fusions neutralize TcdB1 *in vitro* with greater potency than bezlotoxumab

We previously characterized a panel of anti-TcdB Nbs from alpacas immunized with inactive TcdB1 and determined domain specificity and neutralization potential *in vitro* (*22*). An approximate epitope was also determined for a subset of Nbs by negative-stain electron microscopy (nsEM) (*22*). Based on these data, we identified three Nbs with therapeutic potential. Two were specific for different sites on the DD and one for the GTD. We previously denoted these as B0E2, B0D10, and B2C11 but have renamed them here as D1, D2, and G1, respectively. These Nbs were fused to a human IgG1 Fc to form Nb-hFc homodimers with increased half-life and valency (*16, 17*) (Fig. 1A, Table S1). We first tested the neutralization potency of these constructs against TcdB1 in a Vero cell viability assay. While we were unable to generate complete curves for these homodimer Nb-hFc fusions, it was clear that these were less effective than bezlotoxumab (Fig. 1B).

**Figure 1.**
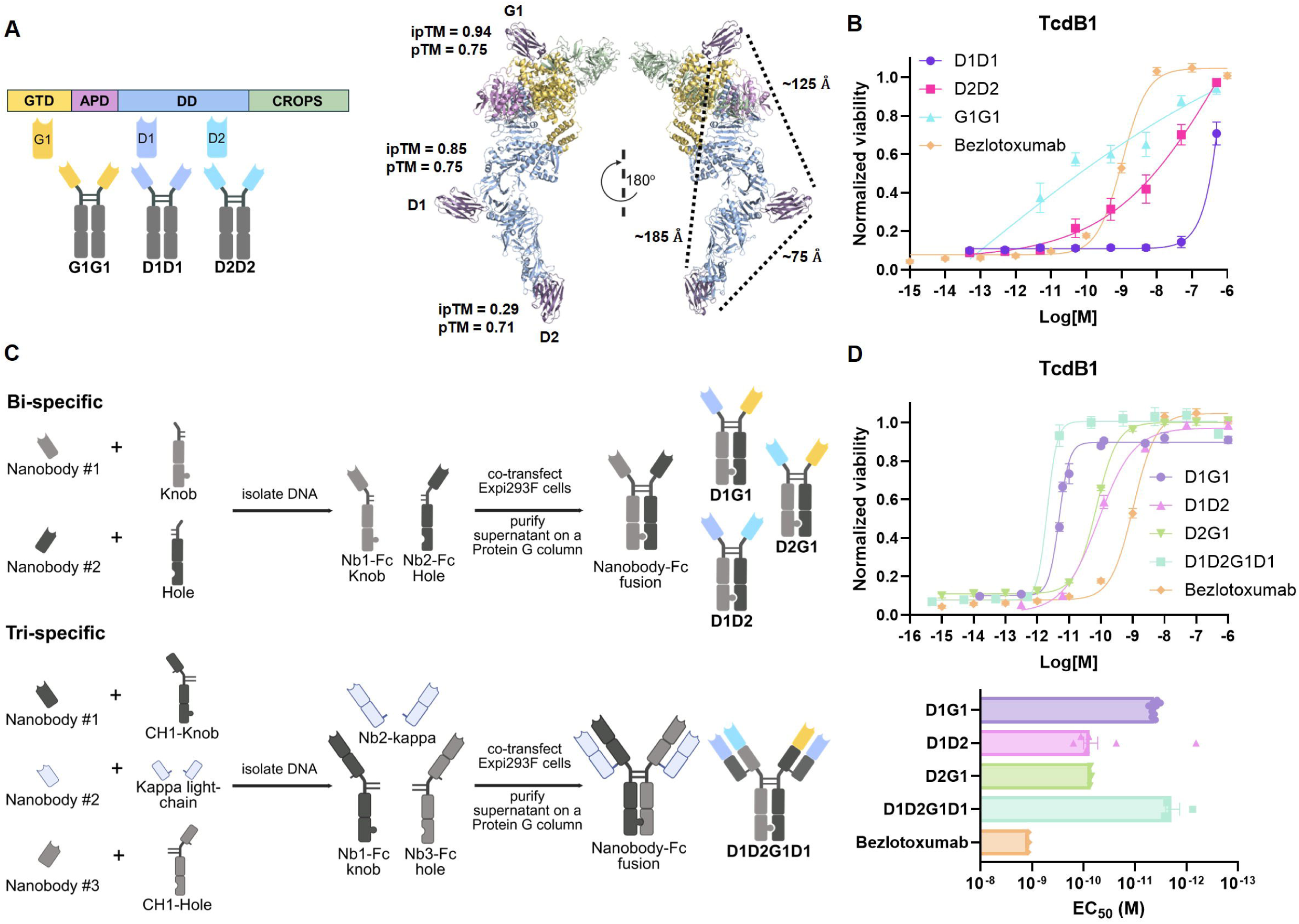
Multi-specific Nb-hFc fusions neutralize TcdB1 with greater potency than homodimer Nb-hFc fusions and bezlotoxumab. **(A)** Domain specificity of three Nbs against TcdB and the resulting homodimer Nb-hFc fusions. Diagram created in BioRender. The model of each Nb with TcdB1 was made in AlphaFold3 (*23*). Distances between Nbs were approximated in PyMOL. **(B)** *In vitro* neutralization of 30 fM TcdB1 with homodimer Nb-hFc fusions, D1D1, D2D2, and G1G1, and bezlotoxumab on Vero cells. Each point represents the mean of three or more biological replicates. Error bars indicate the standard error of the mean. **(C)** Schematic for creating bi- and tri-specific Nb-hFc fusions and the resulting fusions made. Diagram created in BioRender. **(D)** *In vitro* neutralization of 30 fM TcdB1 with multi-specific Nb-hFc fusions, D1G1, D1D2, D2G1, and D1D2G1D1, and bezlotoxumab on Vero cells. Each point represents the mean of three or more biological replicates. Half-maximal effective concentration (EC_50_) values for each fusion and bezlotoxumab were calculated from neutralization curves of individual biological replicates with a full curve (n = 10, 6, 3, 3, 3 for D1G1, D1D2, D2G1, D1D2G1D1, bezlotoxumab, respectively). Error bars indicate the standard error of the mean.

We hypothesized that a bi-specific or tri-specific format could increase neutralization by allowing two sites of TcdB to be simultaneously engaged. We utilized a “knobs-into-holes” approach to generate bi-specific Nb-hFc fusions. This method introduces bulky side chain mutations (“knob”) into the portion of the CH3 domain of an Fc necessary for antibody dimerization, and smaller side chain mutations (“hole”) into the other Fc (*14, 15, 18*). These mutations force heterodimerization and create a bi-specific antibody when the “knob” and “hole” Fc are fused to different variable domains (*18–20*). We generated three bi-specifics, D1G1, D1D2, and D2G1 (Fig. 1C, Table S1). We also created one tri-specific fusion, D1D2G1D1, by adding a CH1 domain, which is necessary for light-chain pairing, to the “knob-Fc” and “hole-Fc” and then fusing the third Nb onto a kappa light-chain constant domain (Fig. 1C, Table S1). Based on AlphaFold3 modeling (*23*) (Fig. 1A), which corroborated our previous nsEM micrographs of the single Nb:TcdB complexes, we expected D1D2 and D1G1 to concurrently access both epitopes while D2G1 would be unable to do so as the D2 and G1 epitopes are on distal ends of the toxin. This was further supported by distance approximations between Nb epitopes in which D1D2 and D1G1 are below the approximate 150 Å maximum distance the IgG flexible hinge provides (*24*), but D2G1 is not. For this reason, we selected D1D2 and D1G1 to constitute each half of the tri-specific fusion. All multi-specifics had greater TcdB1 neutralization in a cell viability assay than bezlotoxumab, with EC_50_ values 10 – 100-fold lower (Fig. 1D).

### Multi-specific Nb-hFc fusions have high affinity and potent neutralization across TcdB subtypes

While sequence similarity is ∼88% or greater for TcdB1/2/3 (*10*), there is the possibility that one of our Nb-hFc fusions could lose affinity for one or more subtypes, as was reported for bezlotoxumab against TcdB2 (*13*). We performed surface plasmon resonance (SPR) to determine the binding kinetics and apparent binding affinities for all homodimer and multi-specific Nb-hFc fusions to TcdB1/2/3. All fusions had sub-nanomolar affinities for TcdB1/2/3, except for D1D1, which lost affinity to TcdB2 (Fig. 2A, Fig. S1, and Table S2). This could imply multi-specific fusions containing D1 may lose neutralization potency against TcdB2. However, the bi- or tri-specific format containing the high affinity Nbs D2 and G1 might overcome D1’s lower affinity to TcdB2 and maintain neutralization potency.

**Figure 2.**
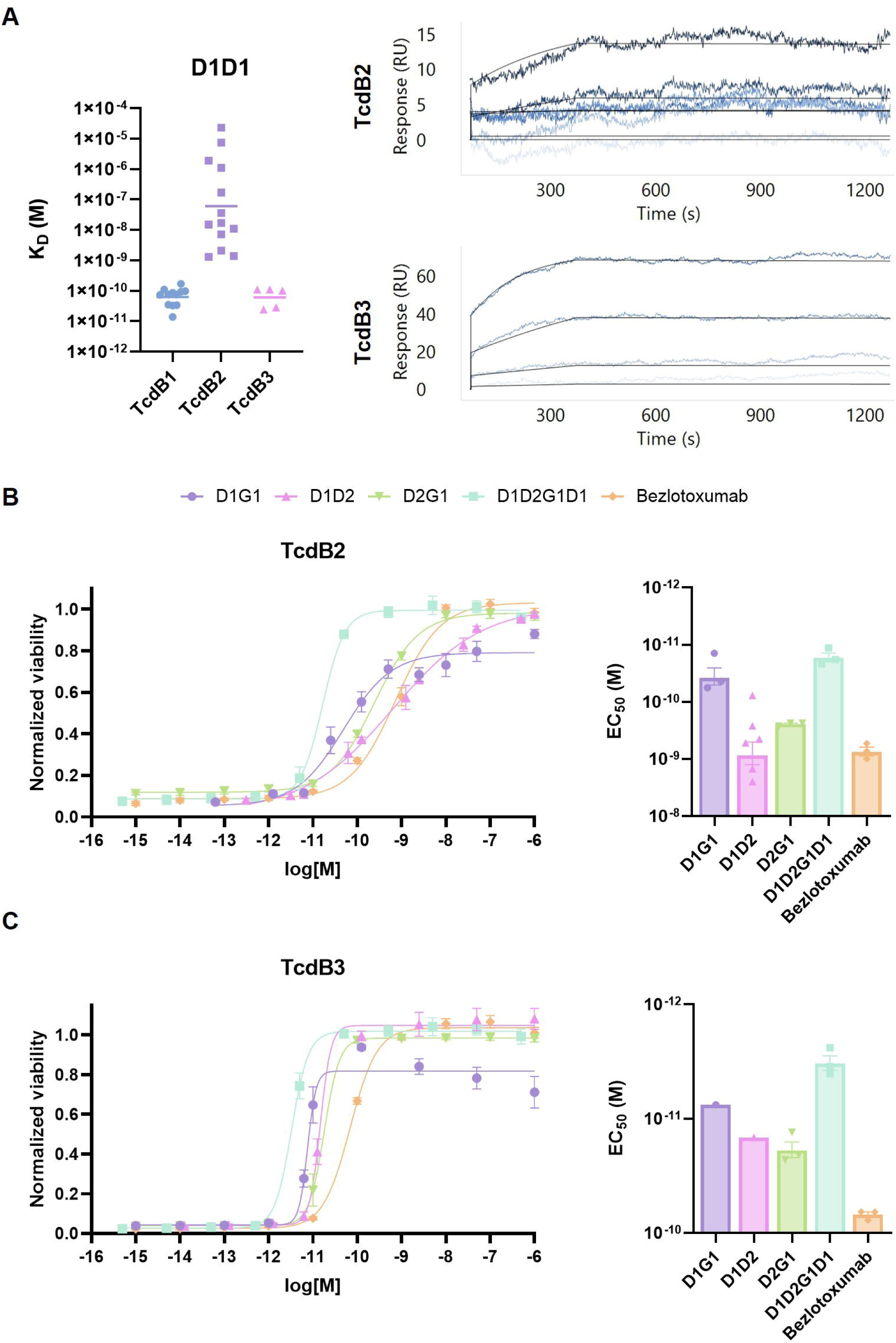
Multi-specific fusions show broad TcdB neutralization despite D1D1 losing affinity to TcdB2. **(A)** Binding affinities of D1D1 for TcdB1/2/3 as determined by SPR with representative SPR curves for D1D1 with TcdB2 and TcdB3. Each data point indicates an individual replicate (n = 12, 11, 5 for TcdB1/2/3, respectively). **(B)** *In vitro* neutralization of 30 fM TcdB2 or **(C)** TcdB3 with multi-specific Nb-hFc fusions, D1G1, D1D2, D2G1, and D1D2G1D1, and bezlotoxumab on Vero cells. Each point represents the mean of three or more biological replicates. Half-maximal effective concentration (EC_50_) values for each fusion and bezlotoxumab were calculated from neutralization curves of individual biological replicates with a full curve (For TcdB2, n = 3 for all groups, except n = 6 for D1D2). For TcdB3, when a fusion did not have any biological replicates with a full curve to calculate an EC_50_, the average curve across biological replicates from panel C was used (n = 3 for all groups, except D1G1 and D1D2 are the average EC_50_ from panel C). Error bars indicate the standard error of the mean.

We further tested neutralization to TcdB2 and TcdB3 to determine if any multi-specific fusions had potential as a pan-neutralizing therapeutic. While all multi-specific Nb-hFc fusions had higher EC_50_ values for TcdB2 than TcdB3, these were still in the picomolar range (Fig. 2B-C). All were more potently neutralizing than bezlotoxumab for both subtypes, as well (Fig. 2B-C). As with TcdB1, the tri-specific fusion had similar neutralization potency as D1G1 against TcdB2 and TcdB3, although the D1G1 data were not ideal, resulting in a poor curve fit (Fig 2B-C). Consistent with the TcdB1 data (Fig. 1B), the neutralization potency of the homodimers was similar to that of bezlotoxumab for TcdB2, except for D1D1, which was non-neutralizing (Fig. S2). This aligned with D1D1’s loss of affinity for TcdB2 (Fig. 2A). Additionally, bezlotoxumab had greater TcdB3 neutralization than all three homodimers (Fig. S2). These experiments highlight the broad and potent neutralization capabilities of all four multi-specific fusions over their homodimer counterparts and bezlotoxumab.

### D1G1, D1D2, and D1D2G1D1 improve infection outcomes against a TcdB2-producing strain

TcdB2 is produced by the NAP1/B1/027 strains of *C. difficile*, which have been associated with epidemic outbreaks (*25, 26*). Since all multi-specific fusions neutralized TcdB2 *in vitro*, we next wanted to test these in a murine model of CDI with the TcdB2-producing strain R20291 (Table S3). We prophylactically treated mice with mAb or Nb-hFc by intraperitoneal injection (IP) 24 hours prior to infection by oral gavage with R20291 spores (Fig. 3A). Along with the four multi-specific fusions, we included a combination therapy of two bi-specific fusions (D1G1 and D1D2) as these fusions make up the two halves of the tri-specific. While all treatment groups lost weight on day 2 post-infection, D1D2 and D1G1 + D1D2 lost less than IgG control pretreated mice (Fig. 3B-C). On day 3 post-infection, all fusions improved weight recovery, except for D2G1 (Fig. 3B-C). D1D2 and D1G1 + D1D2 continued to gain more weight than IgG control and bezlotoxumab pretreated mice on day 4 post-infection (Fig. 3B-C). Interestingly, D1G1 and D1D2G1D1 were the only treatment groups to maintain a 100% survival rate compared to 50% survival in the IgG control group (Fig. 3D). While not significant compared to the IgG control group, D1D2 and D1G1 + D1D2 had a survival rate of 90% (Fig. 3D). Notably, bezlotoxumab and D2G1 did not improve survival relative to the IgG control group (Fig. 3D).

**Figure 3.**
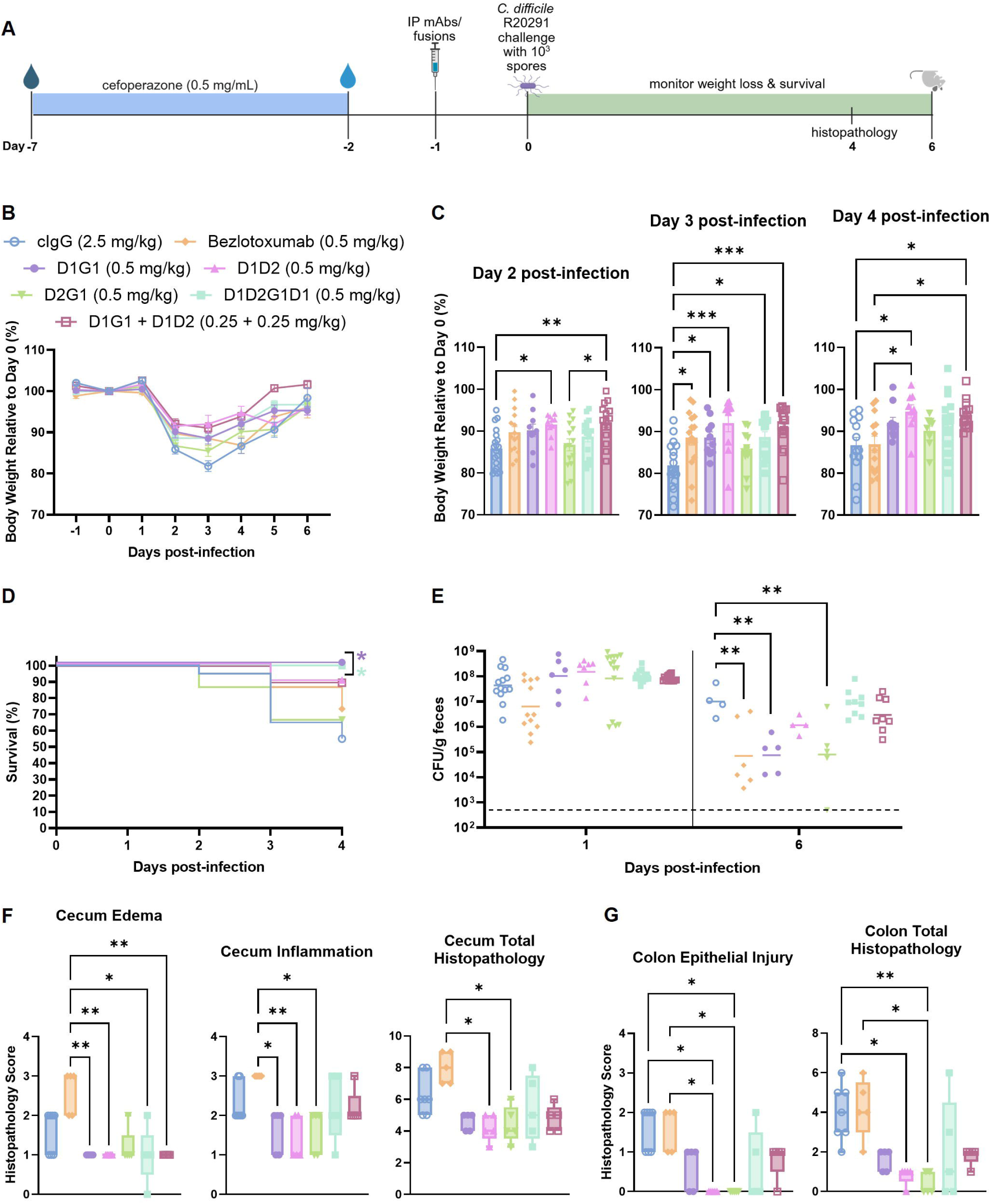
Nb-hFc fusions as prophylaxis against a TcdB2-producing strain, R20291. **(A)** Infection schematic. Diagram created in BioRender. **(B)** Percent body weight relative to baseline in mice pre-treated with D1G1, D1D2, D2G1, D1G1 + D1D2, D1D2G1D1, bezlotoxumab, or control IgG (cIgG) prior to infection with *C. difficile* R20291 (n = 10, 10, 15, 15, 15, 15, 20, respectively). Data points represent group means; error bars indicate the standard error of the mean. **(C)** Individual body weights relative to baseline from panel B on days 2, 3, and 4 post-infection. Data points represent individual mice; the shaded area represents the mean and error bars indicate the standard error of the mean. Data were analyzed by ordinary one-way ANOVA with Tukey’s post-hoc. **(D)** Four-day survival analysis. Curve comparisons were analyzed by logrank (Mantel-Cox) test. **(E)** *C. difficile* total bacterial burden in stool on days 1 and 6 post-infection. Data points represent individual mice and the bar indicates the geometric mean. The dashed line indicates the limit of detection, which is 500 CFUs/g feces. Data were analyzed by lognormal ordinary one-way ANOVA with Dunnett’s post-hoc. **(F)** Selected histopathology scores from Fig. S3 for ceca and **(G)** colons obtained on day 4 post-infection (n = 5 for all treatment groups, except n = 7 for cIgG). Data were analyzed by Kruskal-Wallis test with Dunn’s post-hoc. * P < 0.05, ** P < 0.01, *** P < 0.001.

Bacterial burden was quantified on day 1 post-infection to verify *C. difficile* colonization and on day 6 post-infection to determine if any treatment groups reduced bacterial burden. Initially, all mice were similarly colonized at 10^7^ – 10^8^ colony forming units [CFUs]/g feces (Fig. 3E). However, on day 6 post-infection, D1G1-, D2G1-, and bezlotoxumab-treated mice had reduced colonization to 10^4^ CFUs/g feces, on average (Fig. 3E).

Ceca obtained on day 4 post-infection for all Nb-hFc fusions had no decrease in edema, inflammation, or epithelial injury compared to the control IgG group (Fig. S3A-B). However, there was a decrease in edema for D1G1, D1D2, D1G1 + D1D2, and D1D2G1D1 compared to bezlotoxumab (Fig. 3F). Mice pretreated with D1G1, D1D2, and D2G1 also showed a decrease in cecal inflammation compared to bezlotoxumab-treated mice (Fig. 3F). D2G1 reduced colonic inflammation compared to the IgG control- and bezlotoxumab-treated groups, and both D1D2 and D2G1 showed no scorable epithelial injury, which was significant compared to IgG control and bezlotoxumab mice (Fig. S3C-D, Fig. 3G). D2G1 did not provide any benefit in terms of weight or survival and was omitted from subsequent mouse experiments. Taken together, these results demonstrate that all multi-specific Nb-hFc fusions, excluding D2G1, improved CDI outcomes over bezlotoxumab administered at 0.5 mg/kg.

### D1G1, D1D2, and D1D2G1D1 improve survival against a lethal TcdB1-producing strain

We next wanted to test the Nb-hFc fusions against a lethal TcdB1-producing strain, VPI 10463, using the same prophylactic regimen (Fig. 4A, Table S3). Nb-hFc dosing was increased from 0.5 mg/kg to 2.5 mg/kg to account for the expected increase in disease severity with this strain (*27*). While all treatment groups lost weight on day 2 post-infection, D1D2, D1G1 + D1D2, and D1D2G1D1 prevented the severe weight loss observed in the IgG control and bezlotoxumab pretreated mice (Fig 4B-C). D1G1 + D1D2 and D1D2G1D1 continued to improve weight recovery on day 3 post-infection (Fig. 4C). All IgG control- and bezlotoxumab treated mice succumbed to infection by day 2 and 3 post-infection, respectively (Fig. 4D). All multi-specific fusions significantly increased survival, with D1D2G1D1 at 80%, D1D2 at 70%, and D1G1 and D1G1 + D1D2 at 60% (Fig. 4D). While the multi-specific fusions improved infection outcomes, they did not decrease cecal or colonic histopathology in tissues obtained on day 2 post-infection (Fig. S4). Only D1G1 + D1D2 modestly reduced epithelial injury in the colon in comparison to both the IgG control and bezlotoxumab groups (Fig. S4C-D). CFUs were enumerated over the course of infection with no major differences noted (Fig. S5).

**Figure 4.**
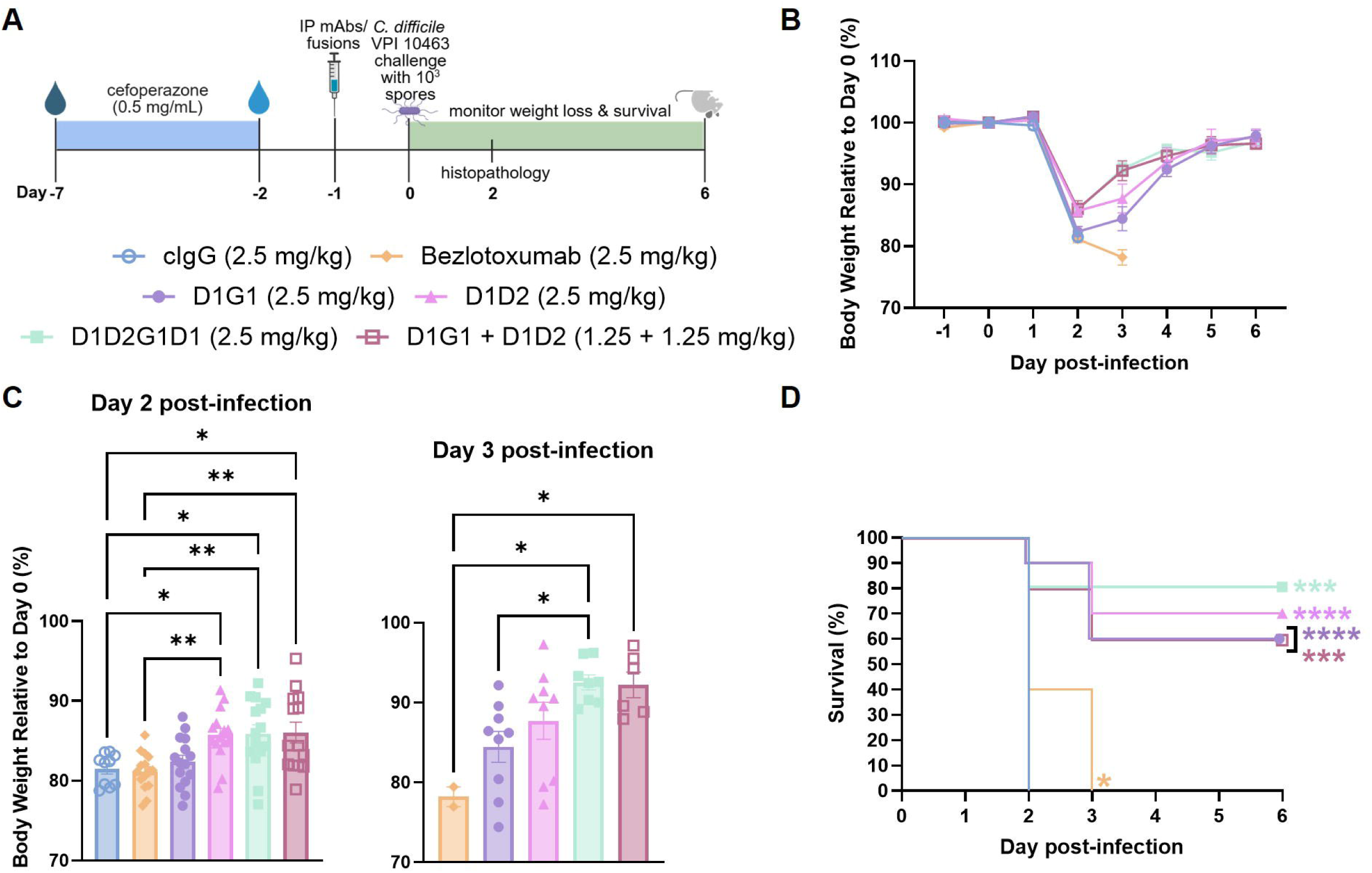
Multi-specific Nb-hFc fusions as prophylaxis against a lethal TcdB1-producing strain, VPI 10463. **(A)** Infection schematic. Diagram created in BioRender. **(B)** Percent body weight relative to baseline in mice pre-treated with D1G1, D1D2, D1G1 + D1D2, D1D2G1D1, bezlotoxumab, or control IgG (cIgG) prior to infection with *C. difficile* VPI 10463 (n = 15 to day 2 post-infection, 5 mice were sacrificed on day 2 post-infection for histology). Data points represent group means; error bars indicate the standard error of the mean. **(C)** Individual body weights relative to baseline from panel B on days 2 and 3 post-infection. Data points represent individual mice; shaded area represents the mean and error bars represent the standard error of the mean. Data were analyzed by ordinary one-way ANOVA with Tukey’s post-hoc. **(D)** Six-day survival analysis. Curve comparisons were analyzed by logrank (Mantel-Cox) test. * P < 0.05, ** P < 0.01, *** P < 0.001, **** P < 0.0001.

### All Nb-hFc fusions reduce cecal edema from infection with a TcdB3-producing strain

To further probe the breadth of protection, we next challenged mice with M68, a TcdB3-producing strain (Fig. 5A, Table S3). Consistent with other reports, this strain did not cause high mortality or morbidity in mice (*27, 28*) (Fig. S6A-B), so clinical outcome was monitored by bacterial burden and tissue pathology. On day 6 post-infection, there was a 10-fold decrease in CFUs for D1G1 and D1G1 + D1D2 relative to the control group (Fig. 5B). Tissues were obtained on day 6 post-infection for blinded histopathology scoring. D1D2 reduced inflammation in the cecum relative to tissue from the IgG control group (Fig. 5C). Although histopathology scores were relatively low for cecal edema for all treatment groups, when visualizing the tissue, there was a distinct difference in edematous area for mice pretreated with IgG control versus those administered the multi-specific fusions or bezlotoxumab (Fig. 5C-D). Quantifying the area of edema showed that all tested multi-specific fusions significantly reduced edema, whereas bezlotoxumab pretreated mice did not (Fig. 5E).

**Figure 5.**
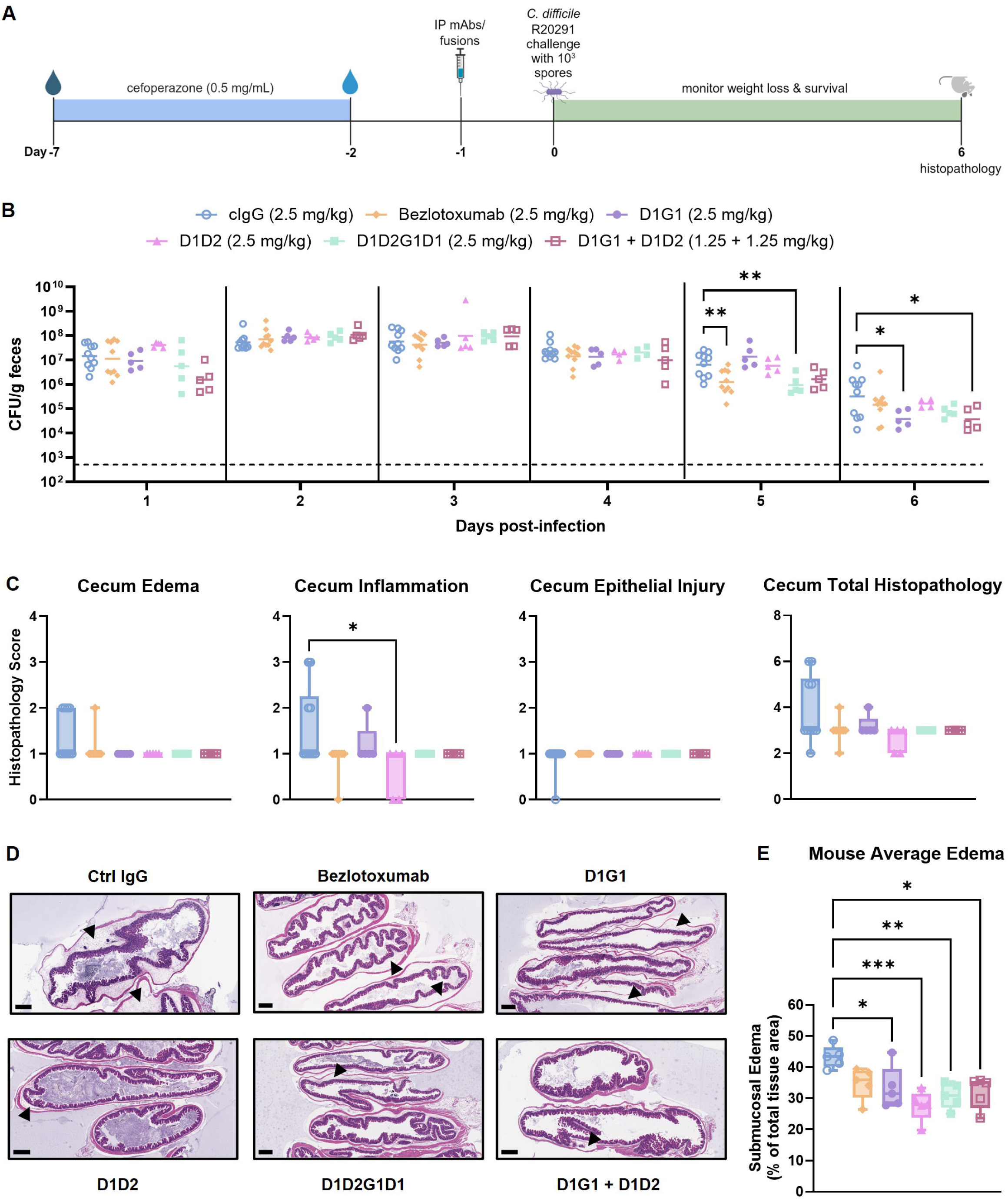
Multi-specific Nb-hFc fusions as prophylaxis against a TcdB3-producing strain, M68. **(A)** Infection schematic. Diagram created in BioRender. **(B)** *C. difficile* total bacterial burden in stool represented as CFU/g feces from days 1 through 6 post-infection. Data points represent individual mice and the bar indicates the geometric mean. The dashed line indicates the limit of detection, which is 500 CFUs/g feces. Data were analyzed by lognormal ordinary one-way ANOVA with Tukey’s post-hoc. **(C)** Blinded histopathological scoring of cecal tissues collected on day 6 post-infection. Data points represent individual mice. Box and whisker plot with min to max values**. (D)** Representative H&E stained cecal sections from each treatment group. Black arrows indicate submucosal edema. Scale bar is 500 µm. **(E)** Quantification of submucosal edema as a percentage of total cecal tissue area. Each data point is the average edema of 2 – 4 cecal sections per mouse. Box and whisker plot with min to max values. * P < 0.05, ** P < 0.01, *** P < 0.001.

### D1G1 improves survival in a therapeutic model of CDI

We finally wanted to determine if there could be therapeutic potential for these multi-specific fusions. As such, we developed a therapeutic model of CDI in which the mice are treated with oral vancomycin and an IP-delivered antibody fusion two days after infection (Fig. 6A), mimicking a canonical CDI relapse/recurrence model (*29, 30*). Instead of monitoring for prevention of relapse, which would require daily cage changes to remove *C. difficile* reservoirs, mice were maintained in their infected cages to ensure relapse and determine if the high mortality observed in this recurrence model could be prevented (*30*). The mice were administered a second dose of the fusions on day 8 post-infection, when vancomycin treatment was ceased, to ensure circulating antibody upon relapse. We prioritized the comparison of one bi-specific fusion, D1G1, to bezlotoxumab against the TcdB2 representative strain. We chose D1G1 as it afforded 100% survival and reduced bacterial burden during acute infection with this strain when given prophylactically (Fig. 3D-E). We observed increased bacterial burden 3 days post-vancomycin cessation beginning on day 11 post-infection and peaking on day 13 post-infection for all treatment groups (Fig. 6B). This increase in bacterial burden corresponded with a drop in weight on day 12 post-infection for IgG control and bezlotoxumab treated mice, whereas treatment with D1G1 significantly prevented weight loss compared to both control and bezlotoxumab treated groups (Fig. 6C). D1G1 treatment appeared to delay weight loss by one day, as all groups did proceed to lose weight on day 13 post-infection (Fig. 6C). Most strikingly, we observed significantly improved survival in mice treated with D1G1 over both the control and bezlotoxumab treated mice (Fig. 6D).

**Figure 6.**
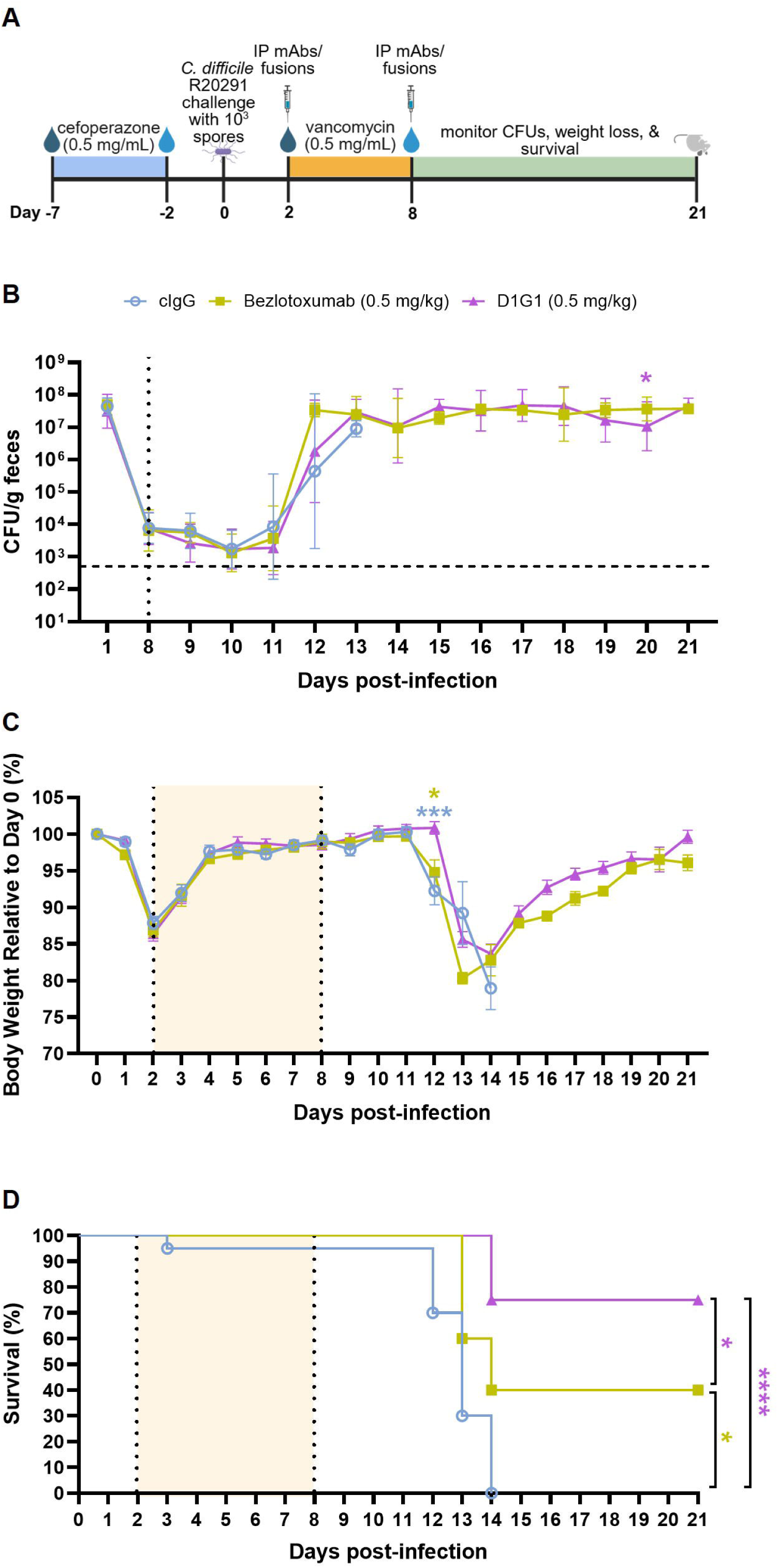
D1G1 protects in a therapeutic model of CDI. **(A)** Infection schematic. Mice were treated with 0.5 mg/kg D1G1, bezlotoxumab, or control IgG (cIgG) on days 2 and 8 post-infection. Diagram created in BioRender. **(B)** *C. difficile* total bacterial burden in stool represented as CFU/g feces. Data points represent the geometric mean; error bars indicate the geometric standard deviation. The dashed line indicates the limit of detection, which is 500 CFUs/g feces. Data on day 20 post-infection were analyzed by one-tailed lognormal Welch’s t-test. **(C)** Percent body weight relative to baseline in mice treated twice with 0.5 mg/kg D1G1, 0.5 mg/kg bezlotoxumab, or cIgG (n = 20, 10, 20 respectively). Data points represent group means; error bars indicate the standard error of the mean. **(D)** 21-day survival analysis. Curve comparisons were analyzed by logrank (Mantel-Cox) test. *P < 0.05, **** P < 0.0001.

## Discussion

Although researchers have developed several antibodies against TcdB, bezlotoxumab was the only biologic with clinical availability; it received FDA approval for the prevention of recurrent CDI in 2016 (*4, 31–35*). The drug’s discontinuation in early 2025 eliminated the only antibody-based treatment option for CDI. There is new appreciation for TcdB sequence variation between strains and there are three TcdB subtypes of high clinical relevance, TcdB1/2/3 (*10, 11*). Given the observation that many antibodies lose affinity and/or neutralization potency to one or more subtypes (*13, 32, 35*), it is important to identify alternative antibodies with broad sequence coverage. We report the generation of three bi-specific and one tri-specific Nb-hFc fusions against TcdB, some of which are more efficacious than the industry standard bezlotoxumab at neutralization of TcdB1/2/3, both *in vitro* and *in vivo*.

Our *in vitro* work demonstrated all multi-specific Nb-hFc fusions had high apparent affinities for TcdB1/2/3 (Fig. S1, Table S2). One exception was the D1D1 homodimer Nb-hFc which had a decreased affinity for TcdB2 (Fig. 2A); we could not accurately assess a binding affinity due to the large variation between replicates. We speculate that this may be because D1 binds near the TcdB1 Frizzled receptor binding site, and the amino acid sequence for this site diverges substantially in TcdB2, resulting in a lower affinity of D1D1 for TcdB2 (*36*). However, the combination of D1 with D2 or G1 in a bi- or tri-specific format was enough to increase the overall affinity for TcdB2. These results highlight how engineered multi-specificity overcomes sequence divergence, a challenge that traditional mAbs cannot address when targeting a single non-conserved binding site. All four multi-specific Nb-hFc fusions, D1G1, D1D2, D2G1, and D1D2G1D1, broadly neutralized TcdB1/2/3 in Vero cell viability assays with potencies exceeding the homodimer Nb-hFc fusions and bezlotoxumab (Fig. 1B&D, Fig. 2B-C, Fig. S2), thus warranting *in vivo* efficacy studies.

We evaluated all multi-specific Nb-hFc fusions and bezlotoxumab as prophylactic treatments in mice infected with representative TcdB1-, TcdB2-, and TcdB3-producing strains. We began our studies with the TcdB2 representative strain, R20291. D2G1 did not provide a weight loss or survival benefit and was removed from further mouse studies (Fig. 3B-D). While not statistically significant, mice pretreated with the remaining fusions, D1G1, D1D2, and D1D2G1D1, showed less mortality than bezlotoxumab-treated mice (Fig. 3D). D1G1 and D1D2 reduced cecal edema and inflammation in mice (Fig. 3F), and D1D2 treated animals also had no colonic epithelial injury (Fig. 3G). These histopathology scores were significantly less than the scores for the bezlotoxumab group, indicating instances of superior tissue protection with D1G1 and D1D2. This is an especially important finding as it is known that bezlotoxumab does not prevent intestinal damage and this provides evidence of superiority for our Nb-hFc fusions (*37*). We did not expect to see a decolonization phenotype, and we did not perform cage changes throughout infection, therefore, the mice are continually exposed to *C. difficile* spores in the cages. However, we did observe a decrease in the fecal bacterial burden for mice pretreated with bezlotoxumab and D1G1 (Fig. 3E). A decrease in bacterial burden is beneficial to reduce the spread of CDI, especially in a hospital setting.

When administered prophylactically against a lethal TcdB1-expresssing strain, D1G1, D1D2, and D1D2G1D1 significantly improved survival, whereas all bezlotoxumab-treated mice succumbed to infection (Fig. 4D). While no fusion completely prevented mortality, dose optimization may further improve survival outcomes. We noted that there were no differences in bacterial burden or histopathology, except for a slight decrease in colonic epithelial injury for the D1G1 + D1D2 combination therapy when compared to the bezlotoxumab group (Fig. S4, Fig. S5). It is possible that the fusions are not reaching the colonic lumen at a high enough concentration before the toxin causes tissue damage by day 2 post-infection. This could be overcome by increasing the treatment dose or frequency.

Since TcdB3-producing strains cause edema rather than mortality (*27*), we quantified cecal edema as a metric of antibody-mediated protection. D1G1, D1D2, and D1D2G1D1 significantly reduced edematous area compared to IgG control-treated mice, whereas the bezlotoxumab-treated group did not (Fig. 5E). Furthermore, D1G1 reduced the bacterial burden in feces as it did against the TcdB2-producing strain, but bezlotoxumab did not as it had previously (Fig. 5B). Taken together, our prophylactic studies demonstrate that D1G1, D1D2, and D1D2G1D1 significantly improve infection outcomes over bezlotoxumab across three diverse strains of *C. difficile*.

Finally, we wanted to determine whether our findings would translate to a therapeutic model with our TcdB2-producing strain. We used D1G1 for this experiment, as this fusion prevented mortality and reduced R20291 colonization. D1G1 treatment delayed the onset of severe infection post-relapse by one day, whereas bezlotoxumab treatment did not, despite both groups having similar bacterial burdens (Fig. 6B-C), thus suggesting that D1G1 can more effectively neutralize TcdB in this model of infection. Additionally, D1G1 significantly improved survival over both IgG control and bezlotoxumab treated mice (Fig. 6D). While we expected to see D1G1 reduce colonization burden as it did in the prophylactic model, this was not observed in this post-antibiotic relapse model. We hypothesize this may be due to further antibiotic treatment with vancomycin after cefoperazone depleting the gut microbiota and allowing for *C. difficile* to persist because of a loss of colonization resistance and decreased competition with other commensal gut bacteria (*29*).

There were several limitations of our work to acknowledge. First, the SPR data should be used for qualitative, but not quantitative comparisons, as a 1:1 fit model was used for the multi-specifics, and both the homodimer and multi-specific hFc fusions showed no measurable dissociation. Second, the Vero cell viability assay showed discrepancies between *in vitro* neutralization and *in vivo* efficacy, specifically for D2G1. We postulate that the effectiveness of D2G1 in neutralizing TcdB activity on cells stems from the concentration of antibody and toxin used in the viability assays, which is likely more than what would be seen during fulminant CDI. While D2G1 may be able to crosslink two toxins *in vitro*, the concentrations are likely insufficient *in vivo*. Further, Vero cells are not a biologically relevant cell line for CDI, but they do contain all the major receptors of TcdB across subtypes, and are standard in the field (*22, 38–40*). Third, for our *in vivo* studies, we did not perform cage changes during infection. This is a confounding variable when evaluating colonization as the mice are coprophagic and may ingest *C. difficile* spores throughout the duration of the experiment. While we highlighted instances where colonization burden was decreased, to determine if there was a notable decolonization phenotype, daily cage changes should be employed. Fourth, we chose not to investigate the possibility of our Nb-hFc fusions to elicit anti-drug antibodies (ADAs) because our Nbs are fused to human IgG1 Fc and not mouse Fc and could elicit some ADAs when investigated in WT C57BL/6J mice. We acknowledge the necessity of future testing to ensure these fusions show minimal to no immunogenicity if they are to be translated as a human therapeutic. Finally, efficacy in a murine model of CDI does not mean it will translate to humans. However, preclinical mouse studies are an important step towards therapeutic translation.

Our study substantiates the therapeutic potential of multi-specific Nb-hFc fusions targeting TcdB for CDI treatment. While other multi-specific anti-TcdB molecules exist (*32, 33*), this is the first report of bi- and tri-specific Nb-hFc fusions that broadly neutralize TcdB1/2/3 *in vitro* and improve infection outcomes *in vivo*. Compared to bezlotoxumab, our fusions exhibited superior potency (lower EC_50_ values), enhanced prophylactic efficacy against representative TcdB1/2/3 strains, and improved therapeutic efficacy against a TcdB2-producing strain. Critically, effective CDI therapeutics must neutralize all clinically relevant TcdB subtypes. Our results provide a foundation for advancing multi-specific broad-spectrum anti-TcdB antibodies into preclinical safety and efficacy studies in larger animal models, and, ultimately, humans.

## Materials and Methods

### Study design

The objective of this study was to identify an antibody-based therapeutic candidate with greater efficacy than bezlotoxumab for the treatment of CDI across diverse strains. We assessed the potential therapeutic use of bi- and tri-specific Nb-hFc fusions rather than a traditional mAb. Bi- and tri-specific Nb-hFc constructs (D1G1, D1D2, D2G1, and D1D2G1D1) were first tested *in vitro* for neutralization potency across TcdB1/2/3 using a Vero cell viability assay, and binding affinities were determined by SPR. All constructs showed broad neutralization which catalyzed further exploration *in vivo* using a murine model of CDI. We conducted prophylactic studies against representative strains for TcdB1/2/3, as well as a therapeutic study with a TcdB2 representative strain. Metrics collected during mouse infection studies included: weight, survival, CFUs, and cecal and colonic tissue. All animal procedures were conducted in strict accordance with our established animal protocol. Mice were humanely euthanized if their weight dropped below 80% of their initial weight or they became moribund. Sample size for mouse studies was determined by power analysis (Δ = 0.5, α = 0.05, β = 0.8). All mice were randomly assigned treatment groups. Histopathology scoring was performed by a board-certified gastrointestinal pathologist blinded to the experimental groups. For all prophylactic mouse studies, the endpoint was predetermined to be 6 days post-infection as no more mortality occurs and mice begin to recover weight. The endpoint for the therapeutic model was not predetermined but rather was ceased once weight recovery was achieved. All *in vitro* work was replicated at least three times, and all *in vivo* work was performed in at least two independent experiments, except the therapeutic model which was performed once.

### Toxin expression and purification

Recombinant TcdB1/2/3 were expressed in *Bacillus megaterium* and purified by a series of Ni-affinity, ion-exchange, and size-exclusion chromatography as previously described (*41–43*). All proteins were sterile-filtered through a 0.22 µm membrane and stored in a 20 mM HEPES (pH 6.9), 50 mM NaCl buffer. Protein concentrations were determined by absorbance using calculated extinction coefficients.

TcdB1: ε_0.1%, 280 nm_, 1.067 M^-1^ cm^-1^; molecular weight, 269,170 Da

TcdB2: ε_0.1%, 280 nm_, 1.085 M^-1^ cm^-1^; molecular weight, 270,505 Da

TcdB3: ε_0.1%, 280 nm_, 1.057 M^-1^ cm^-1^; molecular weight 269,324 Da

### Antibody and human Fc fusion expression and purification

Bezlotoxumab was expressed in Expi293F cells for 7 days prior to harvesting the supernatant. Supernatant was sterile filtered through a 0.22 µm membrane and then loaded onto a protein G column (HiTrap Protein G HP, Cytiva) for purification according to the manufacturer’s instructions.

Multi-specific fusion proteins were expressed and purified the same as for bezlotoxumab, except Expi293F cells were co-transfected with heavy chain “knob” or “hole” plasmid DNA at a 1:1 ratio for bi-specifics and a 1:1:1.5 ratio of heavy chain “knob”:heavy chain “hole”:light chain for the tri-specific.

“Knob” and “hole” mutations are found in the CH3 domain and are S354C, T366W and Y349C, T366S, L368A, Y407V, respectively, where cysteine mutations are for increased stability through disulfide bond formation (*18–20*).

### Cell viability assays with recombinant TcdB1/2/3

Vero cells were seeded at a density of 3,000 cells/well in a 96-well black clear bottom plate or 750 cells/well in a 384-well black clear bottom plate with Eagle’s Minimum Essential Medium (EMEM) supplemented with 10% fetal bovine serum. Cells were incubated overnight at 37 °C with 5% CO_2_. The next day, media on the cells was refreshed. Serial dilutions of mAbs and Nb-hFc fusions were incubated at a 1:1 vol/vol with 0.6 pM TcdB1/2/3 for 30 min at room temperature for antibody:toxin complexes to form. Antibody:toxin serial dilutions were then added to cells at a 1/10 dilution for a total 20-fold dilution from the starting concentration. After 72 hours of incubation, spent media was aspirated and fresh media was added. CellTiter Blue was added per Promega’s protocol, and cells were incubated for an additional 3.5 hours before fluorescence was measured at 560 nm/590 nm excitation/emission by a Cytation 5 plate reader (BioTek). Viability was determined by: (average fluorescence mAb/Nb-hFc/TcdB – average fluorescence media)/(average fluorescence cells – average fluorescence media). EC_50_ values were calculated in GraphPad Prism using the least squares fit of the log(agonist) vs. response -- variable slope (four parameters) model.

### Surface plasmon resonance

Kinetic parameters for the monospecific fusions were determined by surface plasmon resonance using the Carterra LSA^XT^ platform and CDMP biosensor chip (#4282). Multi-specific fusions were also tested, although their kinetic parameters could only be approximated using a 1:1 binding model. The experiment was performed at 25°C using HEPES-buffered saline with Tween (Teknova #H8174) for ligand printing or with the addition of 5 g/L bovine serum albumin as running buffer. The chip was conditioned with 20 mM sodium hydroxide then 10 mM sodium acetate (pH 5), then activated with 2:1:1 0.1 M MES (pH 5.5), 400 mM EDC, and 130 mM NHS, for amine-coupling of anti-human-IgG Fc (Southern BioTech #2104-01) in 10 mM sodium acetate (pH 4.5), and blocked with ethylenediamine in 150 mM borate (pH 8.5). The fusions were printed at 0.625, 1.25, 2.5, and 5 ng/mL, each in quadruplicate for 10 min. Buffer injections and TcdB1/2/3, in increasing 5-fold dilutions from 160 pM to 500 nM, were flowed over the ligands with a 5 min association phase, and 15 min buffer-only dissociation phase. After each TcdB subtype, the surface was regenerated with 0.2 M glycine (pH 2) and ligands were re-printed. Each replicate was curated to exclude ligand or analyte conditions with multivalent behavior, based on high ligand density (>100 response-units) and/or poor fit to a 1:1 Langmuir model. Kinetic parameters were calculated as the geometric mean and 95% confidence interval (geometric mean x/÷ SEM) for replicates of the same or different ligand densities across two independent experiments.

### *C. difficile* spore preparation

Strains were cultured at 37 °C on brain-heart-infusion agar plates supplemented with 0.5% yeast extract, 0.1% cysteine, and 0.1% taurocholate (BHIS-TA) in an anaerobic chamber under these conditions: 90% N_2_, 5% H_2_, and 5% CO_2_. A single colony was then used to inoculate 4 mL BHIS media, which was then incubated overnight at 37 °C under anaerobic conditions. The next day, 2 mL of overnight culture was used to inoculate 40 mL of Clospore sporulation media (1% special peptone mix, 1% yeast extract, 0.06% (NH_4_)_2_S0_4_, 0.012% MgS0_4_ · 7H_2_0, 0.008% CaCl_2_ · 2H_2_0, 0.348% K_2_C0_3_, and 0.26% KH_2_PO_4_) (pH 7.9) and allowed to grow anaerobically for 7 days. Cultures were then centrifuged at 3200 RPM for 20 min at 4 °C, supernatant was discarded, and pellets were washed 3 – 5 times in cold, sterile water. Spore pellets were then resuspended in 1 mL sterile PBS and heat-treated at 55 °C for 20 min to kill vegetative cells. Spore stocks were stored at 4 °C until use.

### Animal studies and experimental design

Female, 8-week old C57BL/6J mice were purchased from Jackson Laboratories (000664) and acclimated to the new facility for at least one week prior to the beginning of any studies, with ad libitum access to chow pellets and water. All studies were performed under the protocol: M1700185-02.

A mouse model of CDI was used for all prophylaxis studies (*44, 45*). 9-week-old C57BL/6J mice were given cefoperazone (0.5 mg/mL) in drinking water, ad libitum, for five days, followed by two days recovery on regular water. 24 hours prior to infection, mice were anesthetized by isoflurane inhalation and injected intraperitoneally with mAbs or Nb-hFc fusions diluted in sterile PBS to a final volume of 100 µl. Mice were infected by oral gavage with 1,000 spores diluted in 100 µL sterile PBS.

For the therapeutic model of CDI, 9-week-old C57BL/6J mice were given cefoperazone (0.5 mg/mL) in drinking water, ad libitum, for five days, followed by two days of recovery on regular water. On day 0 mice were infected by oral gavage with 1,000 spores diluted in 100 µL sterile PBS. On day 2 post-infection, mice were given vancomycin (0.5 mg/mL) in drinking water, ad libitum, for six days. On days 2 and 8 post-infection, mice were intraperitoneally injected with mAb or Nb-hFc fusions diluted in 100 µL sterile PBS.

Mice were monitored daily for weight and signs of morbidity. On specified days, bacterial burden was quantified by macerating stool in sterile PBS and dilution plating on taurocholate-cycloserine-cefoxitin-fructose agar (TCCFA) semi-selective medium comprising 0.1% (w/v) TA, 250 µg/mL D-cycloserine, 16 µg/mL cefoxitin, 0.6% (w/v) fructose, 40 mg/mL Proteose Peptone No. 2, 5 mg/mL Na_2_HPO_4_, 1 mg/mL KH_2_PO_4_, 2 mg/mL NaCl, 0.1 mg/mL MgSO_4_ anhydrous, and 20 mg/mL agar. Mice were humanely euthanized if they lost >20% of their initial body weight or became moribund. All mice were randomly assigned to experimental groups.

### *C. difficile* enumeration

Stool from infected mice was macerated in 1x PBS (pH 7.4) and serially diluted. Serial dilutions were spotted onto TCCFA semi-selective media plates which were grown anaerobically overnight (*46*). After 24 – 48 hours of growth, colony forming units (CFUs) were counted, and *C. difficile* burden, both spore and vegetative, was calculated.

### Histology preparation and tissue scoring

Ceca and colons were harvested post-mortem, flushed with PBS, and fixed in 10% neutral buffered formalin (NBF) at 4°C for 24 hours. Tissues were embedded in paraffin, sectioned (5 µm), and stained with hematoxylin and eosin (H&E). Tissues were scored by a board-certified gastrointestinal pathologist, blinded to treatment groups. Edema, inflammation and epithelial injury were scored based on previously established criteria (*45*). Representative images were captured using an Aperio AT2 whole slide imager. Slides were imaged at 20x magnification with a resolution of 0.5 µm/pixel. Submucosal edema was quantified using image segmentation in QuPath v.0.6.0 as previously described (*27*).

### Statistical analysis

Data were analyzed and graphed using GraphPad Prism v.10.6.1. Parametric tests were used for all analyses, except for ordinal data in which non-parametric analyses were performed. For comparisons between two groups, a one-sided t-test was used. For comparisons between three or more groups, one-way ANOVA was used. Lognormal one-way ANOVA was used when appropriate. Dunnet’s, Tukey’s, and Dunn’s multiple comparisons post-hoc were used where appropriate. Survival curves were analyzed comparing each treatment group to the control IgG group using the log-rank (Mantel-Cox) test. A p-value of less than 0.05 indicated a statistical significance between groups. Missing data points in weight loss graphs for mouse studies are representative of mice succumbing to infection. Missing data points in CFU graphs are representative of mice succumbing to infection or being unable to defecate from the severity of infection.

## Acknowledgments

We thank the entire Lacy lab for their support. We thank the Georgiev Lab at Vanderbilt University Medical Center for providing us with control IgG for all mouse studies. For tissue embedding and H&E staining, we acknowledge the Translational Pathology Shared Resource supported by NCI/NIH Cancer Center Support Grant P30CA068485. Whole slide imaging was performed in the Digital Histology Shared Resource which is supported by the Digestive Disease Research Center and NIH grant P30DK058404.

## Author contributions

Conceptualization: AGE, BWS, DBL

Formal analysis: AGE, BBCL

Funding acquisition: AGE, DBL

Investigation: AGE, BBCL, AJS, AKT, MKW

Methodology: AGE, BBCL, HKK

Supervision: DBL

Visualization: AGE, BBCL

Writing – Original draft: AGE, DBL

Writing – review & editing: AGE, BBCL, HKK, AKT, BWS, DBL

## Data and materials availability

All data are available in the main text or the supplementary materials.

