## Supplemental Data for "Multi-specific nanobody-Fc fusions against TcdB protect *in vivo* from diverse *Clostridioides difficile* strains"

**Supplemental Table S1**. Plasmids used in this study.

| Plasmid # | Construct name | Notes | Source |
|---|---|---|---|
| pBL377 | pC-His1622-*tcdB*1 | pC-HIS1622, $TcdB1_{1-2366}$, strain VPI10463 | Lab stock |
| pBL598 | pC-His1622-*tcdB*2 | pC-HIS1622, $TcdB2_{1-2366}$, strain R20291 | Lab stock |
| pBL1316 | pC-His1622-*tcdB*3 | pC-HIS1622, $TcdB3_{1-2367}$, strain M68 | Lab stock |
| pBL1248 | bezlotoxumab-HC | pCDNA3.4, heavy chain | Lab stock |
| pBL1249 | bezlotoxumab-LC | pCDNA3.4, light chain | Lab stock |
| pNB316 | B2C11-hFc | pCDNA3.4, G1 fused to human Fc | Lab stock |
| pNB312 | B0E2-hFc | pCDNA3.4, D1 fused to human Fc | Lab stock |
| pNB311 | B0D10-hFc | pCDNA3.4, D2 fused to human Fc | Lab stock |
| pNB164 | B0E2-hFc-knobs | pCDNA3.4, D1 fused to human Fc with knobs mutations | Lab stock |
| pNB165 | B2C11-hFc-holes | pCDNA3.4, G1 fused to human Fc with holes mutations | Lab stock |
| pNB317 | B0D10-knobs-hFc | pCDNA3.4, D2 fused to human Fc with knobs mutations | Lab stock |
| pNB308 | B0D10-hFc-holes | pCDNA3.4, D2 fused to human Fc with holes mutations | Lab stock |
| pNB318 | B0E2-kappa-$C_L$ | pCDNA3.4, D1 fused to kappa light chain constant domain | Lab stock |
| pNB319 | B0D10-CH1-knobs-hFc | pCDNA3.4, D2 fused to human Fc with CH1 domain and knobs mutations | Lab stock |
| pNB320 | B2C11-CH1-holes-hFc | pCDNA3.4, G1 fused to human Fc with CH1 domain and holes mutations | Lab stock |

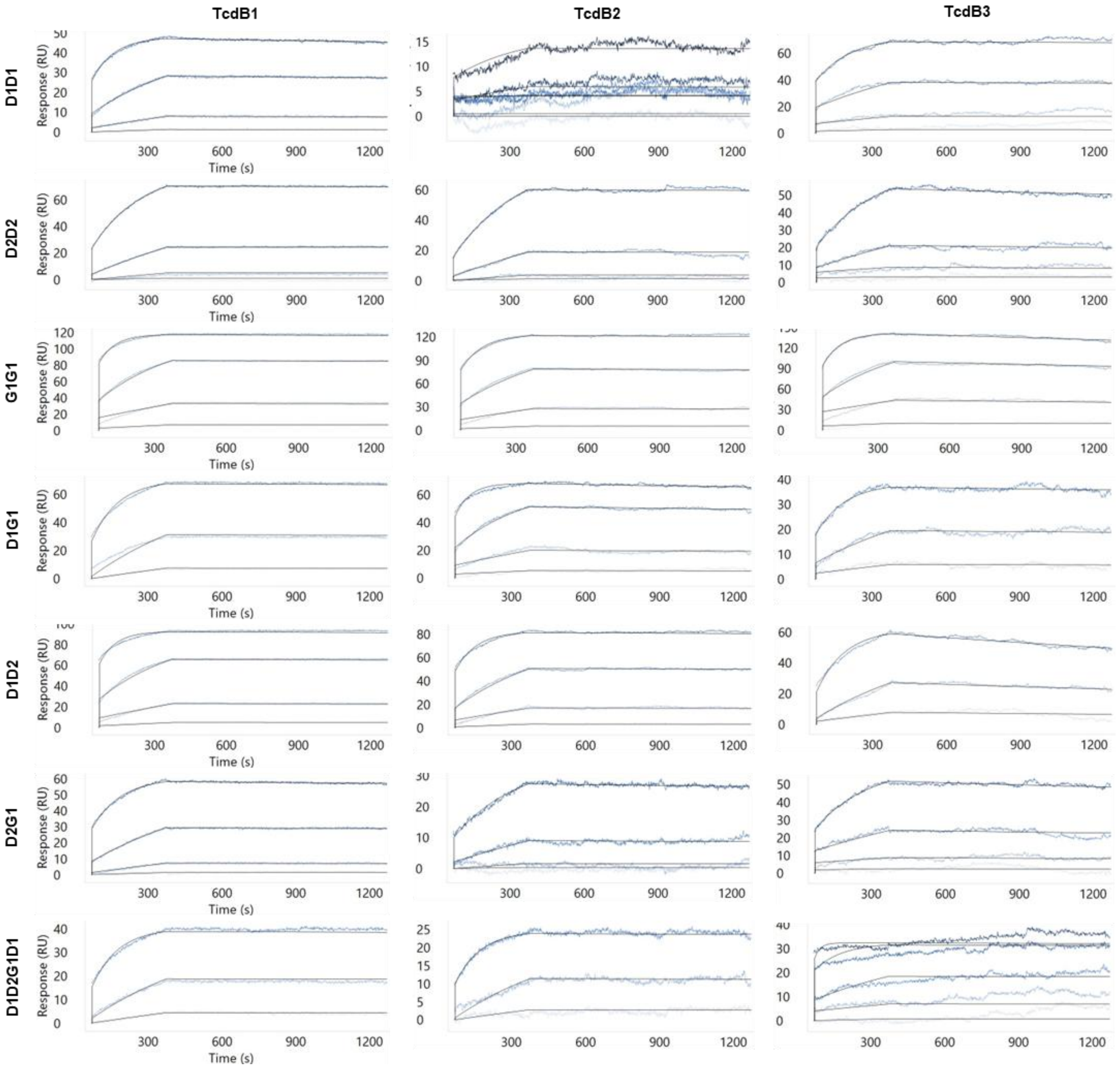


**Supplemental Figure S1. Representative SPR curves for D1D1, D2D2, G1G1, D1G1, D1D2, D2G1, and D1D2G1D1 to TcdB1/2/3.** The fusions were printed at 0.625, 1.25, 2.5, and 5 ng/mL, each in quadruplicate, for 10 min. Buffer injections and TcdB1/2/3, in increasing 5-fold dilutions from 160 pM to 500 nM, were flowed over the ligands with a 5 min association phase, and 15 min buffer-only dissociation phase.

**Supplemental Table S2. Kinetic parameters determined by SPR for all mono- and multi-specific Nb-hFc fusions.** Kinetic parameters were calculated for each fusion against **(A)** TcdB1, **(B)** TcdB2, or **(C)** TcdB3. Geometric mean (GM) and standard error of the mean (SEM) were determined for replicates at the same or different ligand densities, each determined from 3-5 TcdB concentrations. 95% confidence intervals were calculated as (GM x $SEM^{-1}$ – GM x SEM). The number of replicates per fusion varied due to exclusions of conditions with multivalent behavior or, for poor binders, undetectable signal.

| A | TcdB1 | | | |
|---|---|---|---|---|
| | n | $k_a$ ($M^{-1}s^{-1}$ x $10^5$) | $k_d$ ($s^{-1}$ x $10^{-5}$) | $K_D$ (pM) |
| G1G1 | 8 | 11.11 (7.68 – 16.05) | 1.08 (0.98 – 1.24) | 9.70 (7.59 – 12.41) |
| D1D1 | 9 | 6.25 (5.56 – 7.02) | 3.86 (2.46 – 6.06) | 61.89 (37.41 – 102.38) |
| D2D2 | 8 | 2.60 (2.34 – 2.88) | 1.45 (0.89 – 2.35) | 55.59 (33.69 – 91.72) |
| D1G1 | 9 | 17.08 (12.03 – 24.27) | 1.89 (1.15 – 3.10) | 11.16 (5.79 – 21.48) |
| D1D2 | 10 | 9.12 (6.53 – 12.73) | 1.66 (1.03 – 2.69) | 18.07 (8.63 – 37.84) |
| D2G1 | 8 | 4.55 (4.01 – 5.16) | 1.34 (0.90 – 2.01) | 29.55 (18.45 – 47.35) |
| D1D2G1D1 | 8 | 12.17 (7.62 – 19.45) | 2.40 (1.24 – 4.64) | 19.68 (8.29 – 46.73) |

| B | TcdB2 | | | |
|---|---|---|---|---|
| | n | $k_a$ ($M^{-1}s^{-1}$ x $10^5$) | $k_d$ ($s^{-1}$ x $10^{-5}$) | $K_D$ (pM) |
| G1G1 | 8 | 9.19 (5.06 – 16.69) | 2.53 (0.82 – 7.76) | 27.62 (6.29 – 121.29) |
| D1D1 | 3 | 0.003 (0.001 – 0.008) | 21.56 (0.92 – 504.89) | 7.7e5 (1.4e4 – 4.2e7) |
| D2D2 | 8 | 2.37 (1.73 – 3.25) | 1.73 (0.85 – 3.54) | 73.23 (32.42 – 165.39) |
| D1G1 | 4 | 9.25 (8.00 – 10.70) | 4.37 (1.55 – 12.29) | 47.30 (18.90 – 118.40) |
| D1D2 | 7 | 5.09 (4.13 – 6.27) | 2.55 (0.89 – 7.33) | 49.99 (15.44 – 161.87) |
| D2G1 | 3 | 1.96 (1.57 – 2.46) | 1.47 (0.69 – 3.15) | 75.54 (33.07 – 172.54) |
| D1D2G1D1 | 9 | 30.06 (9.47 – 95.35) | 6.57 (1.09 – 39.39) | 21.98 (9.46 – 51.08) |

| C | TcdB3 | | | |
|---|---|---|---|---|
| | n | $k_a$ ($M^{-1}s^{-1}$ x $10^5$) | $k_d$ ($s^{-1}$ x $10^{-5}$) | $K_D$ (pM) |
| G1G1 | 6 | 10.95 (4.32 – 27.72) | 8.08 (1.82 – 35.85) | 73.65 (13.98 – 387.87) |
| D1D1 | 2 | 3.82 (3.36 – 4.34) | 1 (LOD) | < 26.14 (23.02 – 29.70) |
| D2D2 | 6 | 4.47 (1.87 – 10.68) | 2.26 (0.80 – 6.39) | 51.65 (9.29 – 287.10) |

| | | | | |
|---|---|---|---|---|
| **D1G1** | 7 | 52.33 (25.63 – 106.84) | 24.19 (4.39 – 133.38) | 46.31 (7.85 – 273.28) |
| **D1D2** | 8 | 17.79 (9.07 – 34.90) | 25.73 (7.20 – 91.90) | 144.30 (32.70 – 636.39) |
| **D2G1** | 3 | 2.86 (2.21 – 3.70) | 4.70 (0.99 – 22.31) | 163.93 (36.25 – 741.34) |
| **D1D2G1D1** | 2 | 17.69 (0.60 – 517.24) | 16.76 (0.07 – 4208.1) | 95.73 (10.90 – 840.43) |

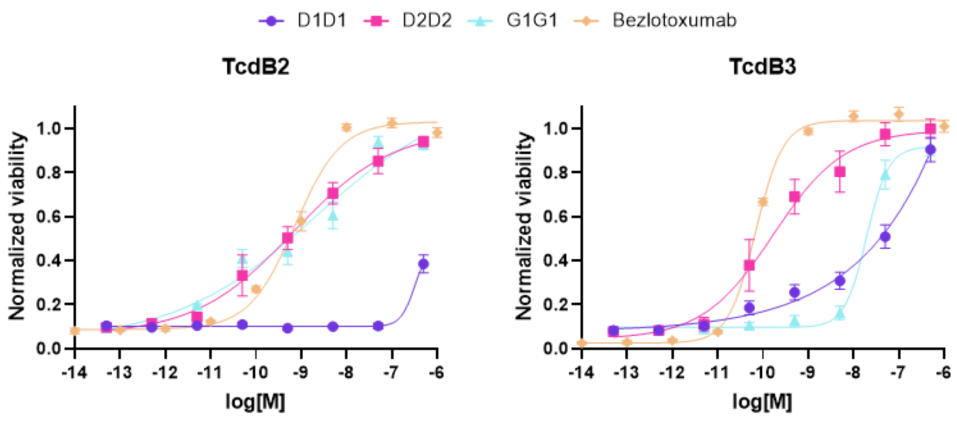


**Supplemental Figure S2. Homodimer Nb-hFc fusions *in vitro* neutralization of TcdB2 and TcdB3 on Vero cells.** Each point represents the mean of three or more biological replicates. Error bars indicate the standard error of the mean.

**Supplemental Table S3.** *Clostridioides difficile* Strains used in this study.

| Strain name | PCR ribotype | Toxins | Source |
|---|---|---|---|
| R20291 | 027 | $TcdA^{+}TcdB2^{+}CDT^{+}$ | Dr. Sarah Kuehne |
| VPI 10463 | 087 | $TcdA^{+}TcdB1^{+}$ $CDT^{-}$ | ATCC |
| M68 | 017 | $TcdA^{-}TcdB3^{+}$ $CDT^{-}$ | Dr. Robert Fagan |

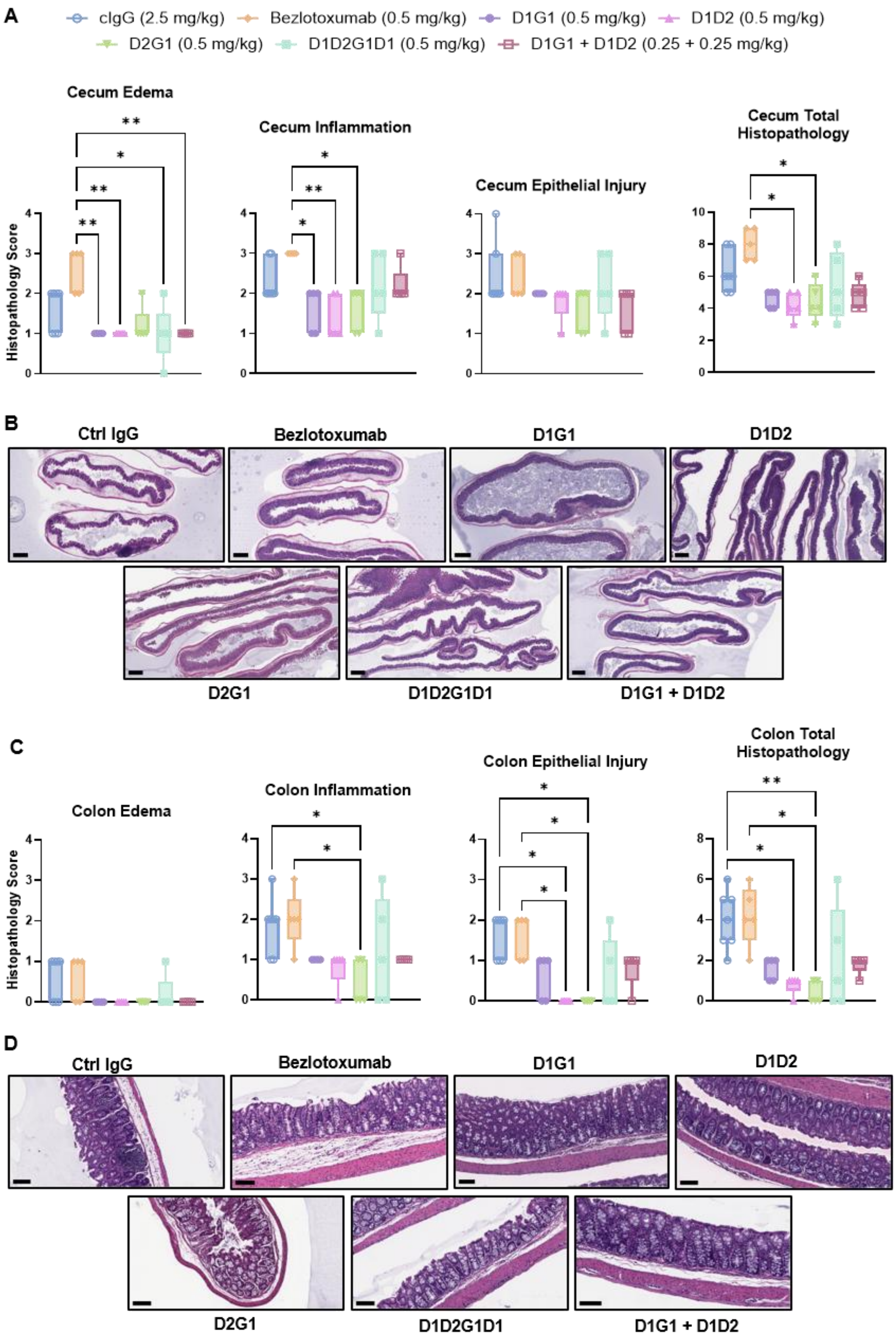
A
cIgG (2.5 mg/kg)
Bezlotoxumab (0.5 mg/kg)
D1G1 (0.5 mg/kg)
D1D2 (0.5 mg/kg)
D2G1 (0.5 mg/kg)
D1D2G1D1 (0.5 mg/kg)
D1G1 + D1D2 (0.25 + 0.25 mg/kg)
Cecum Edema
Cecum Inflammation
Cecum Epithelial Injury
Cecum Total Histopathology
Histopathology Score
B
Ctrl IgG
Bezlotoxumab
D1G1
D1D2
D2G1
D1D2G1D1
D1G1 + D1D2
C
Colon Edema
Colon Inflammation
Colon Epithelial Injury
Colon Total Histopathology
Histopathology Score
D
Ctrl IgG
Bezlotoxumab
D1G1
D1D2
D2G1
D1D2G1D1
D1G1 + D1D2

**Supplemental Figure S3**. **Histopathology scores and representative histology images from R20291 infection. (A)** Histopathology scores for cecal edema, inflammation, and epithelial injury, as well as a summed total histopathology score for tissues obtained on day 4 post-infection (n = 5 for all treatment groups, except n = 7 for the control IgG (cIgG) group). Edema, inflammation, and total histopathology graphs are the same as in Fig. 3F. Data points represent individual mice. Box and whisker plot with min to max values. Data were analyzed by Kruskal-Wallis test with Dunn's post-hoc. **(B)** Representative H&E-stained histology images of the cecum for each treatment group. Scale bar is 500 μm. **(C)** Histopathology scores for colonic edema, inflammation, and epithelial injury, as well as a summed total histopathology score for tissues obtained on day 4 post-infection (n = 5 for all treatment groups, except n = 7 for the cIgG group). Epithelial injury and total histopathology are the same as in Fig. 3G. Data points represent individual mice. Box and whisker plot with min to max values. Data were analyzed by Kruskal-Wallis test with Dunn's post-hoc. **(D)** Representative H&E-stained histology images of the colon. Scale bar is 100 μm.

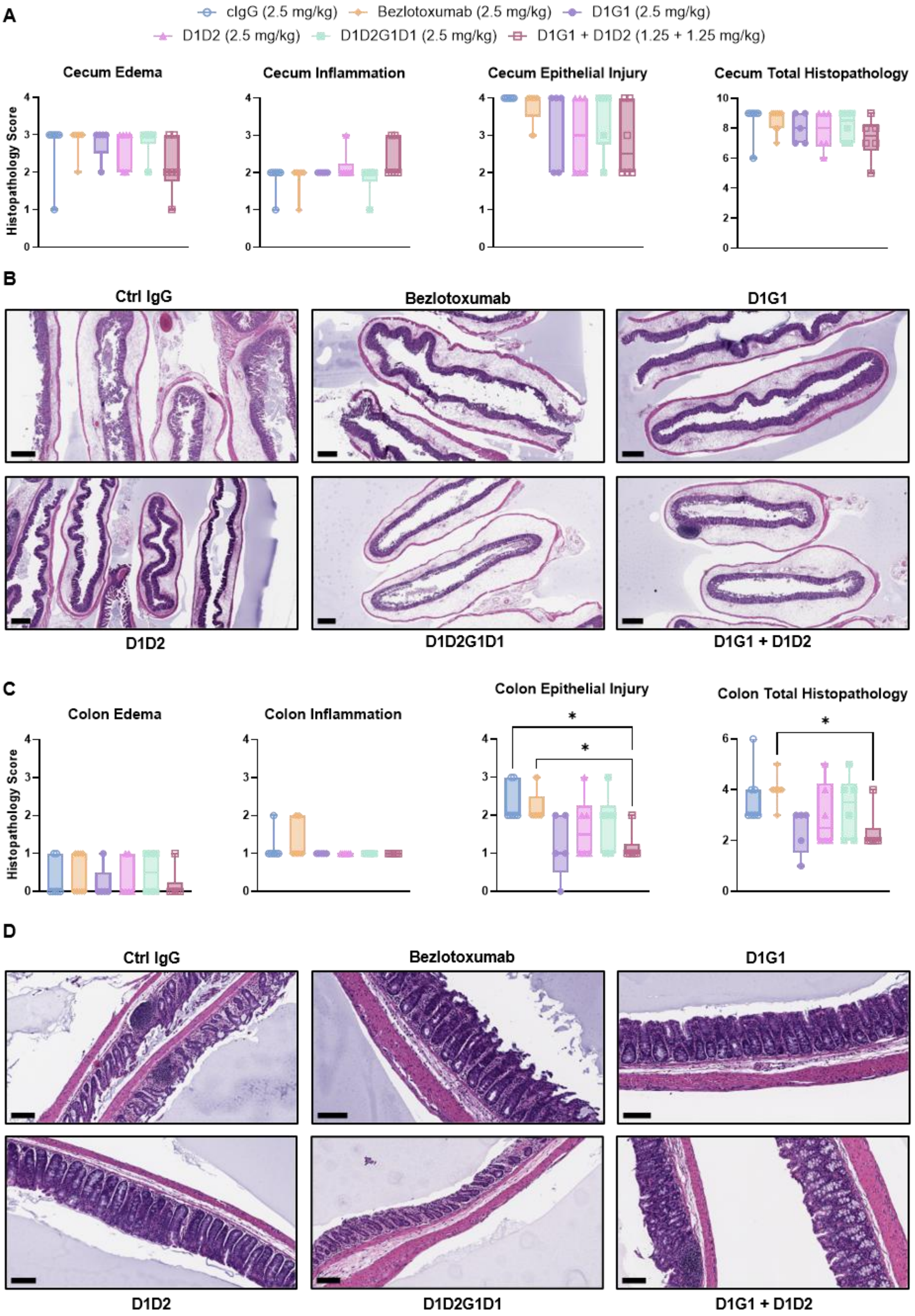
A
cIgG (2.5 mg/kg)
Bezlotoxumab (2.5 mg/kg)
D1G1 (2.5 mg/kg)
D1D2 (2.5 mg/kg)
D1D2G1D1 (2.5 mg/kg)
D1G1 + D1D2 (1.25 + 1.25 mg/kg)
Cecum Edema
Cecum Inflammation
Cecum Epithelial Injury
Cecum Total Histopathology
Histopathology Score
B
Ctrl IgG
Bezlotoxumab
D1G1
D1D2
D1D2G1D1
D1G1 + D1D2
C
Colon Edema
Colon Inflammation
Colon Epithelial Injury
Colon Total Histopathology
Histopathology Score
D
Ctrl IgG
Bezlotoxumab
D1G1
D1D2
D1D2G1D1
D1G1 + D1D2

**Supplemental Figure S4. Histopathology scores and representative histology images from VPI 10463 infection. (A)** Histopathology scores for cecal edema, inflammation, and epithelial injury, as well as a summed total histopathology score for tissues obtained on day 2 post-infection (n = 7, 9, 5, 6, 6, 6, for control IgG (cIgG), bezlotoxumab, D1G1, D1D2, D2G1, D1D2G1D1, respectively). Data points represent individual mice. Box and whisker plot with min to max values. Data were analyzed by Kruskal-Wallis test with Dunn's post-hoc. **(B)** Representative H&E-stained histology images of the cecum. Scale bar is 500 µm. **(C)** Histopathology scores for colonic edema, inflammation, and epithelial injury, as well as a summed total histopathology score for tissues obtained on day 2 post-infection (n = 7, 9, 5, 6, 6, 6, for cIgG, bezlotoxumab, D1G1, D1D2, D2G1, D1D2G1D1, respectively). Data points represent individual mice. Box and whisker plot with min to max values. Data were analyzed by Kruskal-Wallis test with Dunn's post-hoc. **(D)** Representative H&E-stained histology images of the colon. Scale bar is 100 µm.

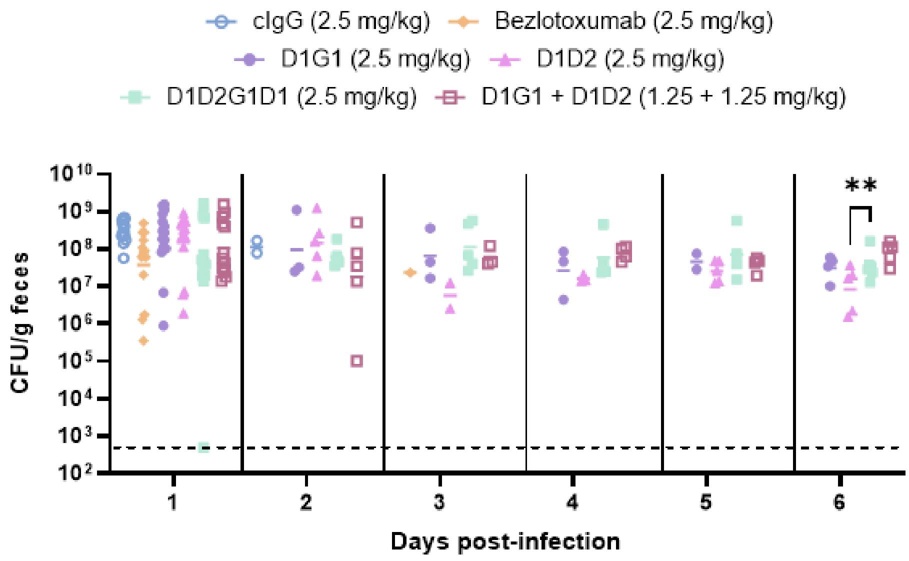


**Supplemental Figure S5.** ***C. difficile*** **burden over the course of infection with VPI 10463.** Data points represent individual mice and the line indicates the geometric mean. The dashed line indicates the limit of detection, which is 500 CFUs/g feces. Data were analyzed by lognormal ordinary one-way ANOVA with Tukey's post-hoc. ** $P < 0.01$.

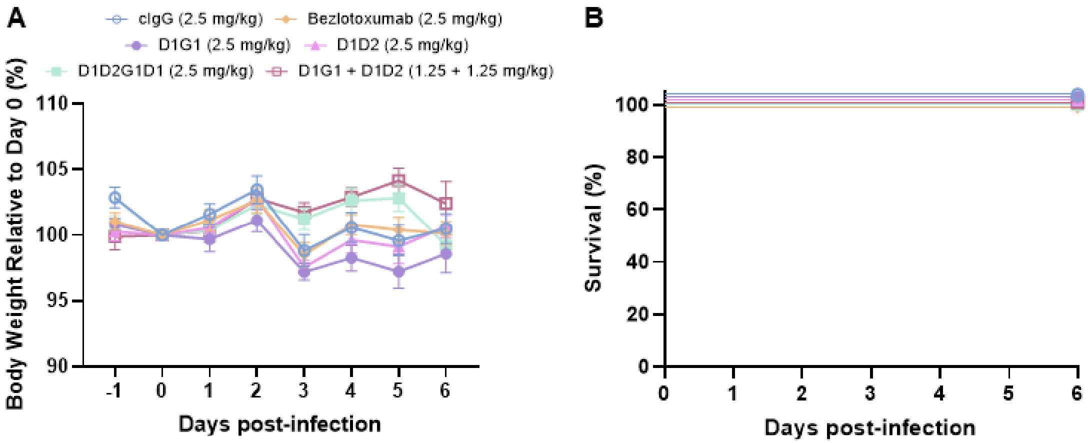


**Supplemental Figure S6.** ***C. difficile*** **strain M68 shows minimal morbidity and no mortality.** **(A)** Percent body weight relative to baseline in mice pre-treated with D1G1, D1D2, D1G1 + D1D2, D1D2G1D1, bezlotoxumab, or control IgG (cIgG) prior to infection with *C. difficile* M68 (n = 5, except n = 10 for bezlotoxumab and cIgG). Data points represent group means; error bars indicate the standard error of the mean. **(B)** Six-day survival analysis. Curve comparisons were analyzed by logrank (Mantel-Cox) test.
